# Two *de novo* transcriptome assemblies and functional annotations from juvenile cuttlefish (*Sepia officinalis*) under various metal and *p*CO2 exposure conditions

**DOI:** 10.1101/2025.06.03.657592

**Authors:** Thomas Sol Dourdin, Antoine Minet, Eric Pante, Thomas Lacoue-Labarthe

## Abstract

The cuttlefish *Sepia officinalis* is a precious model in behavioural and neurobiology studies. It is currently facing combined environmental changes related to the anthropogenic global change. However, genomic resources available to support investigations tackling this issue are still scarce. Therefore, we present two annotated *de novo* transcriptome assemblies from recently hatched (whole body) and one-month-old (head) *Sepia officinalis* juveniles. Both assemblies rely on an important read depth validated by a pseudo-rarefaction analysis, and gathered several individuals from various metal and *p*CO2 exposure conditions. After redundancy reduction, assemblies from newly hatched and one-month-old individuals comprised 230,672 and 370,613 transcripts with 35,590 and 44,233 putative ORFs, respectively, and an annotation rate arounf 70%. Assemblies were compared to each other, revealing age-specific transcriptomic landscapes. These two assemblies constitute highly valuable genomic resources complementing reference genome assembly and facilitating the investigation of transcriptomic endpoints in environmental studies considering coleoid cephalopods.

## Introduction

Cephalopods constitute a remarkable class within molluscs, exhibiting an outstanding diversity of morphologies, habitats and ecology. The class is divided in two subclasses: Nautiloidea Blainville, 1825 and Coleoidea Bather, 1888. While the former contains only a few living fossil species, the latter, counting more than 800 species, represents the large majority of extant cephalopods (*c.a*. octopuses, cuttlefishes, squids). Coleoid cephalopods also display some traits thought to be the result of convergent evolution with vertebrates such as camera-like eye, closed circulatory system and complex central nervous system (Packard, 1972).

So far mostly considered in behaviour and neurobiology studies, cephalopods are becoming promising model systems in ecotoxicology, for biomonitoring (Ajala et al., 2022) as well as experimental studies (García-Flores et al., 2025; Gouveneaux et al., 2023; Penicaud et al., 2017). This growing interest has been supported by the recent development of cephalopod genomic studies (Albertin & Simakov, 2019; Baden et al., 2023) producing several transcriptomes from diverse tissue types (*i.e. Octopus vulgaris* whole body, early life stages (Prado-Álvarez et al., 2022); *Idiosepius pygmaeus* central nervous system and eyes (Thomas et al., 2024)) along with genome assemblies (*i.e. O. vulgaris* (Destanović et al., 2023), *Amphioctopus fangsiao* (Jiang et al., 2022)) during the last years.

Among coleoids, the cuttlefish *Sepia officinalis* is of great economic interest in the northern east Atlantic as one of the main halieutic resource (Arkhipkin et al., 2021). Besides, *S. officinalis* from different areas were shown to be chronically contaminated by various metals such as silver, lead, cadmium or mercury (Ahmed et al., 2022; Ajala et al., 2022; Ihya et al., 2021; Minet et al., 2021; Miramand et al., 2006), as well as anti-depressant compounds like venlafaxine or fluoxetine (Chabenat, Bidel, et al., 2021; Chabenat, Knigge, et al., 2021) leading to behavioural alterations and neurotoxic effects (see Lacoue-Labarthe et al. (2016) for a focus on early-life stages) potentially amplified by ocean acidification (see Minet (2022) for the example of combined exposure with mercury). Yet, despite scientific and ecological relevance, genomic resources for *S. officinalis* remain limited: only one tissue specific transcriptome assembly was released (Gonçalves et al., 2024) and the first chromosome-level reference genome is about to be published (Rencken et al., 2025). High-quality transcriptomes provide essential complement to genome assemblies (Elliott, 2014; Martin & Wang, 2011) by capturing tissue- and condition-specific expression patterns, as well as providing qualitative diversity of isoforms resulting from alternative splicing or RNA editing – particularly abundant in cephalopods and promoting proteome plasticity (Rosenthal & Eisenberg, 2025; Voss & Rosenthal, 2023).

In this context, we present two *de novo* transcriptome assemblies built from the whole body of newly hatched juveniles (project “rnasep1”) and head tissues of one-month-old juveniles (project “rnasep2”) *S. officinalis* reared in various metal and *p*CO_2_ conditions. The gathering of individuals from different experimental conditions enabled the provision of valuable ready-made genomic resources, facilitating future gene expression analyses and comparative transcriptomic studies in coleoid cephalopods.

## Methods

### Description of the biological material

Freshly laid eggs of common cuttlefish (*Sepia officinalis*) were collected in the Minimes Port, Bay of Biscay, France (46°08’44.7”N; 1°10’16.4”W) during the Spring of 2016 (rnasep1) and 2021 (rnasep2). For both experiments, eggs were gathered from multiple independent clutches, and each experimental replicate contained eggs originating from different parental pairs, thereby minimizing potential clutch-or family-specific effects.

### Newly hatched juveniles

Upon field collection, eggs were immediately transported to the laboratory. They were individualized from their clutches and then placed in eighteen 9 L glass aquaria (n=30 eggs per aquarium; salinity: 34.4 ppt; temperature: 18.4 ± 0.6°C; 12 h light:12 h dark cycle, with aeration, recirculation pumps). After 5 days of acclimation, nine aquaria were maintained at ambient seawater pH (pHT = 7.95 ± 0.03) and the nine others were gradually acidified by bubbling CO_2_ until reaching a pHT of 7.70 ± 0.08. The *p*CO_2_ regulation was handled by an IKS system from Aquastar© and weekly calibration of IKS probes with NBS standard pH solutions (pH 4.0 and 7.0), a glass electrode and using TRIS buffer solutions (salinity 35, provided by A. Dickson, Scripps Institution of Oceanography, San Diego, USA). Among each pH condition, two set of three aquaria were spiked with AgNO_3_ or HgCl_2_ solutions in order to reach 1 µg.L^-1^ of metal in seawater, respectively. The last three aquaria were kept metal-free for control conditions. This resulted in 6 rnasep1 experimental conditions: Control (hereon “CT”), acidification (“*p*CO_2_”), exposed to HgCl_2_ (“HgCl_2_”), exposed to AgNO_3_ (“AgNO_3_”), exposed to both acidification and HgCl_2_ (“*p*CO_2_ + HgCl_2_”) or exposed to acidification and AgNO_3_ (“*p*CO_2_+ AgNO_3_”) (Figure 1a). Water renewal was done every two days and spikes of metals were renewed accordingly to maintain the metal concentrations as constant as possible.

**Figure 1.**
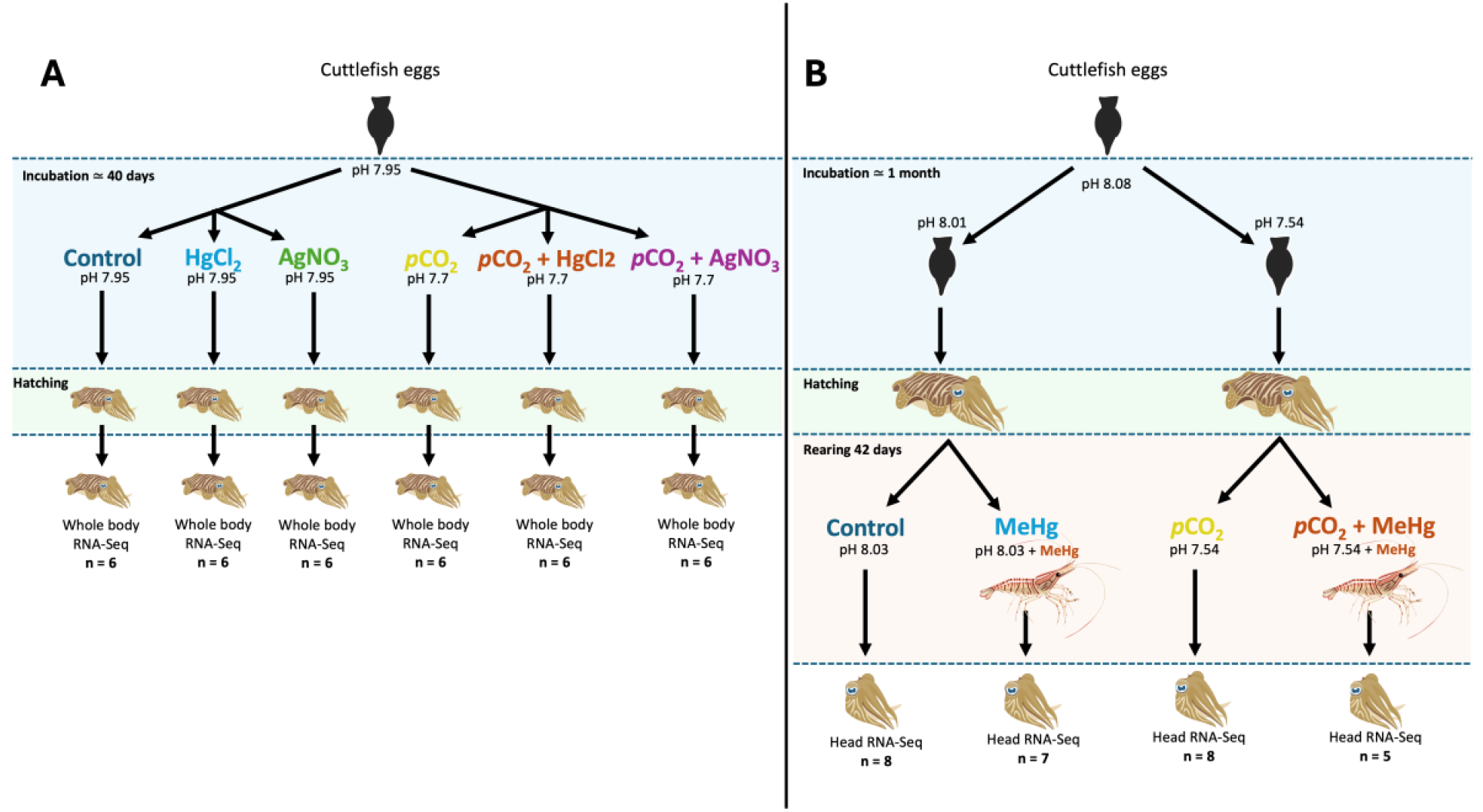
Experimental design for A) newly hatched (“rnasep1”; carried out in 2016) and B) one-month-old juveniles (“rnasep2”; carried out in 2021) exposure to heavy metals and hypercapnia.

Eggs were maintained in these conditions for 40 days, until hatching. When hatched (considering a daily survey), juveniles were immediately collected, anaesthetized with 17.5 mg.L-1 MgCl_2_ and eventually put to death using 2% ethanol in seawater. Then, individuals (n = 6 per condition) were frozen in liquid nitrogen and stored at -80°C before RNA extraction that was performed straight away. Additionally, fourteen hatchlings were sampled for measurement of Ag and Hg concentrations in whole body to validate the exposure conditions (data available in sample description at ENA PRJEB89715, as well as in table S1a).

### One-month-old juveniles

Eggs were acclimated in the laboratory and raised until one-month old juvenile stage to investigate the effects of *p*CO_2_ and MeHg on behaviour of cuttlefish early life stages (Minet, 2022). Briefly, eggs were individualized from their clutches and then acclimated for two weeks in four black plastic basins (*i.e*. batches) in control conditions (n=70 eggs per batch; salinity: 35.0 ppt; temperature: 18.3 ± 0.2°C; pH_NBS_: 8.0 ± 0.1; 12 h light:12 h dark cycle, with aeration, recirculation pumps and daily water renewal). After the acclimation period, two batches remained in control conditions (pH_NBS_: 8.03 ± 0.05 in 2020 and 8.01 ± 0.05 in 2021, equivalent to ∼440 µatm), while the two others were gradually acclimated to acidified conditions by reducing pH_NBS_ (CO_2_ bubbling) from 8.01 to 7.54 ± 0.06 (equivalent to ∼1600 µatm). The *p*CO_2_ regulation was handled by an IKS system from Aquastar© (pH measurements every 20 min and weekly calibration of IKS probes with NBS standard pH 4.0 and 7.0 solutions and a glass electrode.

Immediately after hatching, 27 individuals from each pH condition were transferred into six 20 L black plastic basins and individualized in cylindrical baskets of 60 mm width (*c.a*. 9 indiv. per batch in triplicate). Thereafter, juvenile cuttlefishes were fed *ad libitum* with control (half of the individuals) or MeHg-contaminated (other half of the individuals) live ditch shrimp (*Palaemon varians*). For the MeHg-contaminated condition, only the first shrimp of each day was contaminated to ensure the number of contaminated shrimps eaten to be the same for all MeHg-exposed individuals. This experiment resulted in 4 distinct conditions: Control (hereon “CT”), Acidification (“*p*CO_2_”), contaminated by methyl-mercury (“MeHg”) or a combination of acidification and methyl-mercury contamination (“*p*CO_2_+MeHg”) (Figure 1b).

Hatchlings were raised for 42 days before being anaesthetized with 17.5 mg.L-1 MgCl_2_ and eventually sacrificed using 2% ethanol in seawater. In order to investigate the transcriptomic responses in the whole central nervous system to the experimental conditions, the heads of half of the individuals (CT: n=8; *p*CO_2_: n=8; MeHg: n=7; *p*CO_2_+MeHg: n=5) were immediately sampled and freeze dried in liquid nitrogen before storage at -80°C until RNA extraction. Total mercury concentrations were measured in the muscle of mantle to validate the dietary exposure (data available in sample description at ENA PRJEB88225, as well as in Table S1b).

The experiments were performed in accordance with the French and the European regulations (Directive 2010/63/EU) for the use of animals for scientific purposes. The experimental procedures were referenced and authorized by the French Ministry of Higher Education and Research for the use of animals for scientific purposes (APAFIS # 20520-2019050614554709).

### RNA extraction and Illumina sequencing

### Newly hatched juveniles

RNA extraction was performed using the Nucleospin for NucleoZOL (Macherey Nagel©), using manufacturer instructions. Whole individuals were first ground in liquid nitrogen. The resulting power was incubated in NucleoZOL + RNase-free water (>700 µL/50 mg of tissue powder) for 15 min after 15 s of vigorous vortexing. A 700 µL aliquot was then used for downstream RNA extraction steps. RNA was eluted in 60 µL RNase-free H_2_O. A 5 µL aliquot was sampled for quantification on Qubit (using Thermo Fisher Scientific© HS and BR quantification kits) and 1% gel electrophoresis; the rest of the RNA extraction was immediately placed at -80°C until next steps. Further quality control on a Fragment Analyser (Agilent©), library construction and Illumina sequencing were outsourced to GeT-PlaGe platform from GenoToul, using the Illumina TruSeq Stranded mRNA Library Prep kit and an Illumina HiSeq 3000 platform (2x150bp).

### One-month-old juveniles

The RNA extraction and library preparation were outsourced to Fasteris©. Briefly, 28 heads underwent manual grinding in liquid nitrogen and shredded tissues were then preserved in RLT and beta-mercaptoethanol. Total RNA was isolated using the RNeasy 96 kit (Qiagen©, ref 74181) with final elution in 50 µL of nuclease-free water. After the extraction step, samples were quantified by Quant-iT (Thermo Fisher Scientific©) and the quality of RNA was checked by loading 2 µL of a dilution on a Fragment Analyzer (Agilent©). All samples reached the required concentration of 250-500 ng in a total volume of 50 µL. After poly(A) tail selection, libraries were constructed using the Illumina TruSeq Stranded mRNA Library Prep kit with NEXTFLEX unique dual index barcodes (Revvity©). The libraries were sequenced on an Illumina NovaSeq 6000 platform (2x150bp).

### Pre-assembly processing

The same pipeline of bioinformatic analyses (Figure 2) was carried out for rnasep1 and rnasep2, using the IFB-core high-performance computing cluster of the Institut Français de Bioinformatique (IFB) (ANR-11-INBS-0013). The raw data quality was assessed using FastQC (fastqc_v0.12.1) (Andrews, 2010) in combination with MultiQC (multiqc_v1.14) (Ewels et al., 2016). Fastp (fastp_v0.23.1) (Chen, 2023) was used to filter out adapter sequences as well as low-quality reads (PHRED score ≤ 30), and retrieve only post-filtering reads longer than 120 bp. Kraken2 (kraken2_v2.14) was used to identify and remove any prokaryote (bacteria and archaea) contamination.

**Figure 2.**
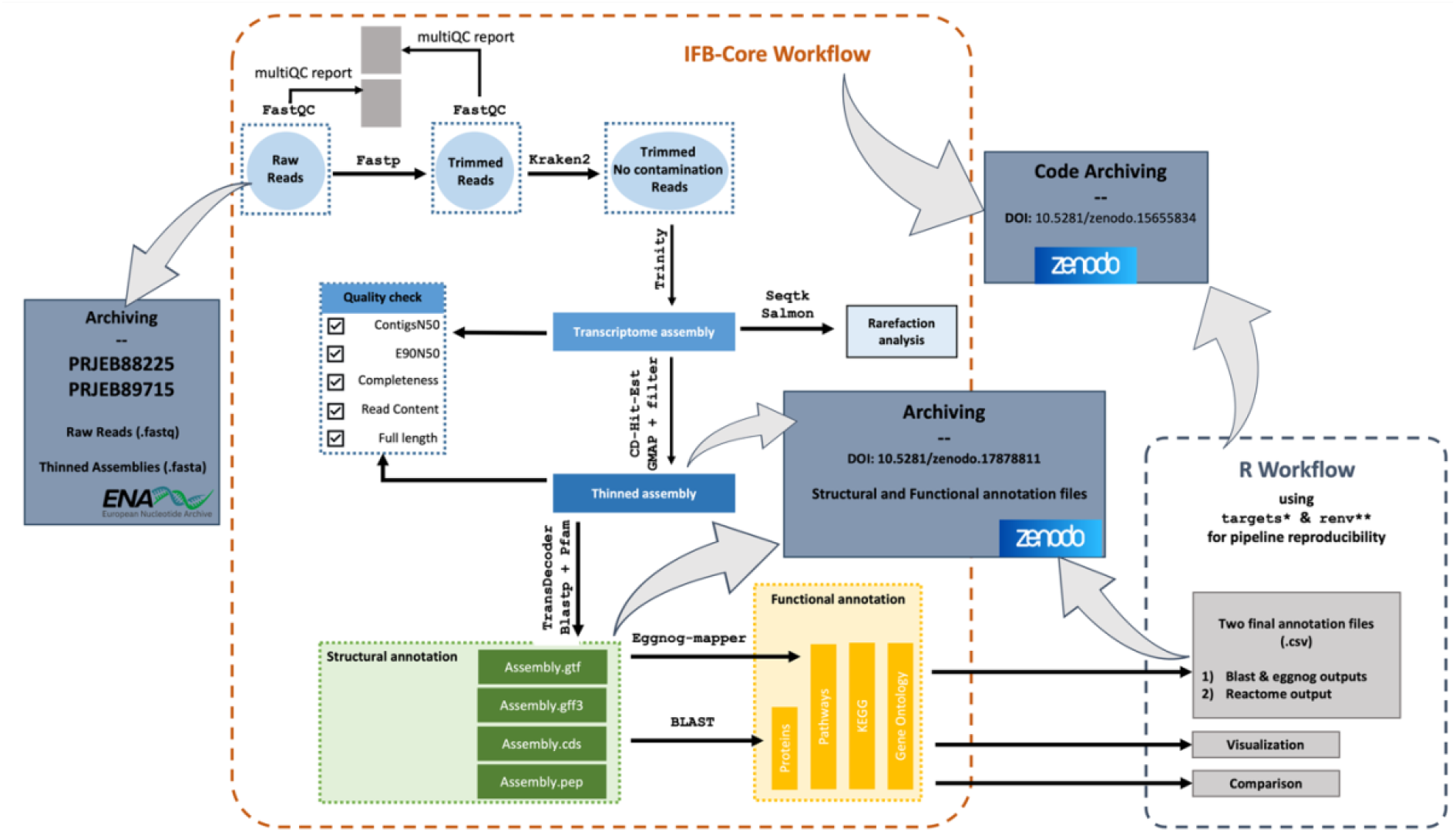
Bioinformatic workflow from raw data to final assembly and annotation files. ^*^(Landau, 2021); ^**^(Ushey & Wickham, 2025)

### *De novo* transcriptome assembly and thinning

Eighteen samples (n=3 per condition) and 16 samples (n=4 per condition) were randomly selected to be included in the rnasep1 and rnasep2 assemblies, respectively, to reduce the time and resource consuming. The *de novo* transcriptome assembly was performed using Trinity (trinity_v2.15.1) (Grabherr et al., 2011) with default parameters and *in-silico* normalization.

Assembled isoforms were clustered based on their similarities (default percentage identity threshold used: 95%) using the CD-HIT tool (cd-hit_v4.8.1) (Li & Godzik, 2006) to retain only the most representative sequences, thereby reducing transcriptome redundancy likely to result from the Trinity method. The resulting assemblies were mapped onto the genomic sequence GCA_050097725.1 (Rencken et al., 2025) using GMAP (gmap_v2020.06.01) (Wu & Watanabe, 2005). From the gff3 file produced, we identified, then filtered out chimeras transcripts (mapping on multiple chromosomes or opposite strands, with too long introns and rearranged exons), as well as transcripts with alignments < 100 bp or < 20% of their total length. Transcripts smaller than 200 bp were removed as well to comply with the European Nucleotide Archive standards.

### Assessment of the transcriptome assemblies

Several metrics were used to fully assess the quality of the assembly. First, TransRate (transrate_v1.0.3) (Smith-Unna et al., 2016) retrieved general statistics from the raw and thinned assemblies. Then, the contigs N50 statistic (at least 50% of the assembled transcript nucleotides are found in contigs that are at least of N50 length), based on the longest isoform per gene and the E90N50 statistic (N50 for top 90% most highly expressed genes) were computed using dedicated Trinity utilities and Kallisto (kallisto_v0.46.2) (Bray et al., 2016). BUSCO (Busco_v5.5.0) (Manni et al., 2021) was used to evaluate the completeness of the transcriptome against two ortholog databases: Metazoa (metazoan_odb10) and Mollusca (mollusca_odb10). The full-length transcript analysis provided by Trinity was also performed using BLAST (blast_v2.16.0) (Camacho et al., 2009) and STAR (star_v2.7.11a) (Dobin et al., 2013) was used to align back the reads against the assembly to assess its read content.

To confirm that the reduced number of samples still ensure a sufficient read depth to catch all available transcripts, we performed an *ad hoc* rarefaction analysis as follows. For each assembly, all the input R1 and R2 fastq files were concatenated, then reads were sub-sampled (keeping pair information) using seqtk (seqtk_v1.3). (https://github.com/lh3/seqtk). For newly-hatched individuals, 1 to 80 million read pairs were sub-sampled. For one-month-old juveniles, 1 to 300 million read pairs were sub-sampled. Then, Salmon (salmon_v1.10.2) (Patro et al., 2017) was used to index Trinity assemblies and quantify the number of reads from each corresponding sub-sample that matches each Trinity transcript. Finally, the number of Trinity transcripts matched by at least 1 read was reported for each sub-sample. (Haas, n.d.) (Camacho et al., 2009)

### Transcriptome annotation

Transcriptome annotation was performed stepwise. First, the putative coding regions (open reading frames [ORFs] ≥ 100 amino acids) were identified by Transdecoder (transdecoder_v5.7.0) (Haas, n.d.), and the prediction was enhanced by homology evidence from BLASTp (blast_v2.16.0) (Camacho et al., 2009) and domain detection using Pfam (Mistry et al., 2021) *via* HMMER/hmmscan (hmmer_v3.3.2) (Eddy, 2011). To generate a representative protein set for downstream annotation, we retained only the longest ORF for each transcript isoform. Then, predicted protein-coding sequences were annotated using a two-step BLASTp strategy. We first performed similarity searches against a curated molluscan UniProt/SwissProt (Bateman et al., 2023) database to obtain taxonomically relevant annotations. Sequences with no significant hits were subsequently queried against the full UniProt/SwissProt database to obtain additional functional annotations. The two BLASTp steps were run using an e-value threshold of 1e-5, retaining only the best hit per query (Bateman et al., 2023). The best hit was retained for each query and its associated Gene Ontology (GO) terms were extracted from the UniProt GO annotation file (*goa_uniprot_all.gaf.gz*). EggNOG-mapper v2 (eggnog-mapper_v2.1.12) (Cantalapiedra et al., 2021) tool was used to complement homology-based annotations by performing functional annotations based on precomputed orthologous groups and complete the Blast annotation with hits from the PFAM (Mistry et al., 2021), COG (Tatusov et al., 2000), KEGG (Kanehisa et al., 2017) and GO (Ashburner et al., 2000) databases. Searches were performed using DIAMOND, restricting the taxonomic scope to bilateria (NCBI TaxID 33213) and retaining only the one-to-one orthologs with an e-value cutoff of 0.001.

### Comparison between newly hatched and one-month-old juveniles

The predicted protein sequences from both assemblies were compared to each other through a reciprocal BLASTp (blast_v2.16.0) (Camacho et al., 2009) to identify reciprocal best hits (RBH) considered to be proteins shared between both assemblies, as well as sequences exhibiting no hit considered as putatively specific. The number of RBH and no-RBH sequences between the two assemblies was represented using the ggvenn R package (ggvenn_v0.1.19). The running of GO enrichment analysis on the lists of specific sequences using GO_MWU script (Fisher’s exact test, FDR: 0.1) did not identified significant enrichment. Therefore, the top 20 specific GO terms from ‘biological_process’ namespace were displayed in scatter plots and summarised by their level 6 ontology using the ontologyIndex (ontologyindex_v2.12) (Greene et al., 2017) R package to facilitate interpretation.

## Results

### Illumina sequencing and pre-processing

The Illumina sequencing produced a total of 358,400,00 paired-end reads for newly hatched and 830,584,708 paired-end reads for one-month-old juveniles (see Tables 1 and 2 for sequencing statistics). The pre-processing resulted in 350,000,000 and 1,108,000,000 cleaned reads accounting for 48.8% and 66.7% of the raw data for newly hatched and one-month-old juveniles, respectively. Then, taxonomic classification of reads did not identify any prokaryote contamination.

**Table 1.** Sequencing statistics for newly hatched juveniles. GeT-PlaGe sample code is used for the labelling of reads in the European Nucleotide Archive repository. R1 GC% and R2 GC% are percent GC content on the forward and reverse reads, respectively.

| GeT-PlaGe Sample Code | Sample Name | Number of Paired-end Reads | R1 GC% | R2 GC% |
| --- | --- | --- | --- | --- |
| Ag771 | SEP1-Ag7.7_1 | 21,283,196 | 40 | 41 |
| Ag8-1-102 | SEP1-Ag8.1_102 | 21,080,270 | 41 | 41 |
| Ag7-7-29 | SEP1-Ag7.7_29 | 17,316,330 | 41 | 41 |
| Ag772 | SEP1-Ag7.7_2 | 19,177,260 | 41 | 41 |
| Ag7-7-31 | SEP1-Ag7.7_31 | 22,963,224 | 41 | 41 |
| Ag7-7-34 | SEP1-Ag7.7_34 | 20,399,022 | 41 | 41 |
| Ag773 | SEP1-Ag7.7_3 | 17,805,326 | 41 | 41 |
| Ag8-1-21 | SEP1-Ag8.1_21 | 23,068,454 | 41 | 41 |
| Ag8-1-27 | SEP1-Ag8.1_27 | 21,202,072 | 40 | 41 |
| Ag812 | SEP1-Ag8.1_2 | 19,480,720 | 41 | 41 |
| Ag813 | SEP1-Ag8.1_3 | 20,526,026 | 41 | 41 |
| Ag816 | SEP1-Ag8.1_6 | 21,223,080 | 41 | 41 |
| Hg7-7-125 | SEP1-Hg7.7_125 | 24,256,850 | 41 | 41 |
| Hg771 | SEP1-Hg7.7_1 | 22,357,978 | 40 | 41 |
| Hg772 | SEP1-Hg7.7_2 | 19,811,618 | 40 | 41 |
| Hg7-7-48 | SEP1-Hg7.7_48 | 20,937,382 | 41 | 41 |
| Hg7-7-49 | SEP1-Hg7.7_49 | 19,721,682 | 41 | 41 |
| Hg7-7-52 | SEP1-Hg7.7_52 | 19,410,228 | 41 | 41 |
| Hg8-1-111 | SEP1-Hg8.1_111 | 21,700,826 | 41 | 42 |
| Hg811 | SEP1-Hg8.1_1 | 17,635,966 | 40 | 41 |
| Hg812 | SEP1-Hg8.1_2 | 24,195,958 | 40 | 40 |
| Hg8-1-35 | SEP1-Hg8.1_35 | 20,576,074 | 41 | 41 |
| Hg8-1-36 | SEP1-Hg8.1_36 | 19,962,954 | 40 | 40 |
| Hg816 | SEP1-Hg8.1_6 | 20,235,544 | 41 | 41 |
| CT7-7-15 | SEP1-CT7.7_15 | 18,380,114 | 40 | 41 |
| CT7-7-18 | SEP1-CT7.7_18 | 18,430,516 | 41 | 42 |
| CT772 | SEP1-CT7.7_2 | 17,710,208 | 40 | 40 |
| CT773 | SEP1-CT7.7_3 | 16,961,648 | 40 | 40 |
| CT774 | SEP1-CT7.7_4 | 15,762,346 | 40 | 40 |
| CT7-7-93 | SEP1-CT7.7_93 | 18,755,500 | 41 | 41 |
| CT811 | SEP1-CT8.1_1 | 20,027,126 | 40 | 41 |
| CT812 | SEP1-CT8.1_2 | 18,261,584 | 41 | 41 |
| CT813 | SEP1-CT8.1_3 | 18,859,762 | 41 | 41 |
| CT8-1-7 | SEP1-CT8.1_7 | 19,050,674 | 41 | 41 |
| CT8-1-82 | SEP1-CT8.1_82 | 20,522,436 | 41 | 41 |
| CT8-1-9 | SEP1-CT8.1_9 | 17,506,642 | 40 | 41 |

**Table 2.** Sequencing statistics for one-month-old juveniles. Fasteris sample code is used for the labelling of reads in the European Nucleotide Archive repository. R1 GC% and R2 GC% are percent GC content on the forward and reverse reads, respectively.

| Fasteris Sample Code | Sample Name | Number of Paired-end Reads | R1 GC% | R2 GC% |
| --- | --- | --- | --- | --- |
| AVRX-1 | SEP2-CT8.1_5 | 31,819,911 | 42 | 42 |
| AVRX-2 | SEP2-CT8.1_6 | 37,005,211 | 43 | 43 |
| AVRX-3 | SEP2-CT8.1_8 | 29,634,371 | 42 | 43 |
| AVRX-4 | SEP2-CT8.1_9 | 33,148,390 | 42 | 43 |
| AVRX-5 | SEP2-CT8.1_37 | 25,239,430 | 42 | 42 |
| AVRX-6 | SEP2-CT8.1_38 | 31,105,749 | 43 | 43 |
| AVRX-7 | SEP2-CT8.1_39 | 33,593,329 | 41 | 41 |
| AVRX-8 | SEP2-CT8.1_80 | 28,294,978 | 42 | 43 |
| AVRX-9 | SEP2-CT7.7_10 | 29,927,066 | 43 | 43 |
| AVRX-10 | SEP2-CT7.7_11 | 27,023,641 | 43 | 43 |
| AVRX-11 | SEP2-CT7.7_12 | 31,302,866 | 42 | 42 |
| AVRX-12 | SEP2-CT7.7_13 | 26,791,740 | 43 | 44 |
| AVRX-13 | SEP2-CT7.7_46 | 33,106,734 | 42 | 42 |
| AVRX-14 | SEP2-CT7.7_48 | 27,834,839 | 45 | 46 |
| AVRX-15 | SEP2-CT7.7_85 | 27,648,758 | 43 | 43 |
| AVRX-16 | SEP2-CT7.7_86 | 30,032,397 | 43 | 43 |
| AVRX-17 | SEP2-Hg8.1_19 | 29,059,690 | 43 | 43 |
| AVRX-18 | SEP2-Hg8.1_20 | 32,381,027 | 44 | 44 |
| AVRX-19 | SEP2-Hg8.1_21 | 24,445,510 | 43 | 45 |
| AVRX-20 | SEP2-Hg8.1_24 | 30,206,664 | 42 | 43 |
| AVRX-21 | SEP2-Hg8.1_25 | 31,415,272 | 43 | 43 |
| AVRX-22 | SEP2-Hg8.1_55 | 24,646,853 | 43 | 43 |
| AVRX-23 | SEP2-Hg8.1_56 | 30,177,602 | 42 | 42 |
| AVRX-24 | SEP2-Hg7.7_28 | 28,964,924 | 42 | 45 |
| AVRX-25 | SEP2-Hg7.7_30 | 39,767,058 | 42 | 41 |
| AVRX-26 | SEP2-Hg7.7_31 | 26,843,136 | 41 | 42 |
| AVRX-27 | SEP2-Hg7.7_32 | 44,104,080 | 41 | 41 |
| AVRX-28 | SEP2-Hg7.7_68 | 25,479,196 | 41 | 41 |

### *De novo* transcriptome assembly quality assessment

For newly hatched juveniles, the thinned transcriptome assembly contained 230,572 transcripts with an average length of 677 bp (Table 3). The quality of the assembly was validated thanks to several metrics. About 83% and 95% of the orthologs from the BUSCO Mollusca and Metazoa databases respectively matched the assembly. In addition, 1,301 proteins matched Trinity transcripts from 80% to 90% of their protein length and 4,892 proteins were represented by nearly full-length transcripts. Besides, 91,3% of the reads used in the assembly process mapped back to the transcriptome. A more detailed alignment report is in Table S2a.

**Table 3.** Transcriptome assembly and assessment metrics for Raw and Thinned assemblies for newly hatched juveniles.

| Type | Metric | Raw | Thin |
| --- | --- | --- | --- |
| Baseline |  |  |  |
|  | Number of Trinity "genes" | 207,672 | 196,944 |
|  | Number of transcripts | 282,660 | 230,572 |
|  | GC content (%) | 36.4 | 36.01 |
|  | Smallest transcript | 174 | 200 |
|  | Largest transcript | 36,551 | 36,551 |
|  | Mean transcript length | 741 | 677 |
|  | Bases with N (ambiguity) | 0 | 0 |
|  | Proportion of N bases | 0 | 0 |
|  | N50 (longest isoform per gene) | 758 | 752 |
|  | E90N50 | 1,932 | 1,864 |
|  | TransRate predicted number of transcripts with ORF | 35,681 | 24,804 |
|  | Mean ORF (%) | 49.43 | 47.99 |
| Completeness |  |  |  |
|  | Mollusca_odb Complete and Single-copy (%) | 43.9 | 53.8 |
|  | Mollusca_odb Complete and duplicated (%) | 43.0 | 28.8 |
|  | Metazoa_odb Complete and Single-copy (%) | 61.7 | 72.1 |
|  | Metazoa_odb Complete and duplicated (%) | 36.2 | 23.1 |
| FullLength |  |  |  |
|  | Number of proteins matching Trinity transcript from 80% to 90% of their protein length | 1295 | 1,301 |
|  | Number of proteins represented by nearly full-length (>80%) transcripts | 4901 | 4,892 |
| ReadContent |  |  |  |
|  | Aligned reads (%) | 97.8 | 91.3 |

The one-month-old juveniles transcriptome assembly obtained after redundancy reduction comprised 370,613 transcripts with a mean length of 569 bp (Table 4). A high proportion of conserved orthologs from the BUSCO Mollusca (83%) and Metazoa (95%) databases were found in the assembly. Moreover, 1,352 proteins matched Trinity transcripts from 80% to 90% of their protein length and 4,927 proteins were represented by nearly full-length transcripts. Finally, 93.4% of the reads used to build the assembly mapped back to it. A more detailed alignment report is in Table S2b.

**Table 4.** Transcriptome assembly and assessment metrics for Raw and Thinned assemblies for one-month-old juveniles.

| Type | Metric | Raw | Thin |
| --- | --- | --- | --- |
| Baseline |  |  |  |
|  | Number of Trinity "genes" | 331,430 | 311,810 |
|  | Number of transcripts | 483,699 | 370,613 |
|  | GC content (%) | 36.94 | 36.49 |
|  | Smallest transcript | 179 | 200 |
|  | Largest transcript | 33,040 | 32,845 |
|  | Mean transcript length | 621 | 569 |
|  | Bases with N (ambiguity) | 0 | 0 |
|  | Proportion of N bases | 0 | 0 |
|  | N50 (longest isoform per gene) | 594 | 591 |
|  | E90N50 | 2,005 | 1,977 |
|  | TransRate predicted number of transcripts with ORF | 43,409 | 28,159 |
|  | Mean ORF (%) | 49.5 | 48.5 |
| Completeness |  |  |  |
|  | Mollusca_odb Complete and Single-copy (%) | 38.9 | 50.5 |
|  | Mollusca_odb Complete and duplicated (%) | 48.6 | 32.3 |
|  | Metazoa_odb Complete and Single-copy (%) | 59.5 | 70.1 |
|  | Metazoa_odb Complete and duplicated (%) | 38.8 | 24.9 |
| FullLength |  |  |  |
|  | Number of proteins matching Trinity transcript from 80% to 90% of their protein length | 1467 | 1352 |
|  | Number of proteins represented by nearly full-length (>80%) transcripts | 5388 | 4927 |
| ReadContent |  |  |  |
|  | Aligned reads (%) | 98.6 | 93.4 |

Rarefaction analysis showed that, for both dataset, the subset of individuals used for assembly is an acceptable compromise between exhaustiveness of transcript inventory and computational effort (Figure 3).

**Figure 3.**
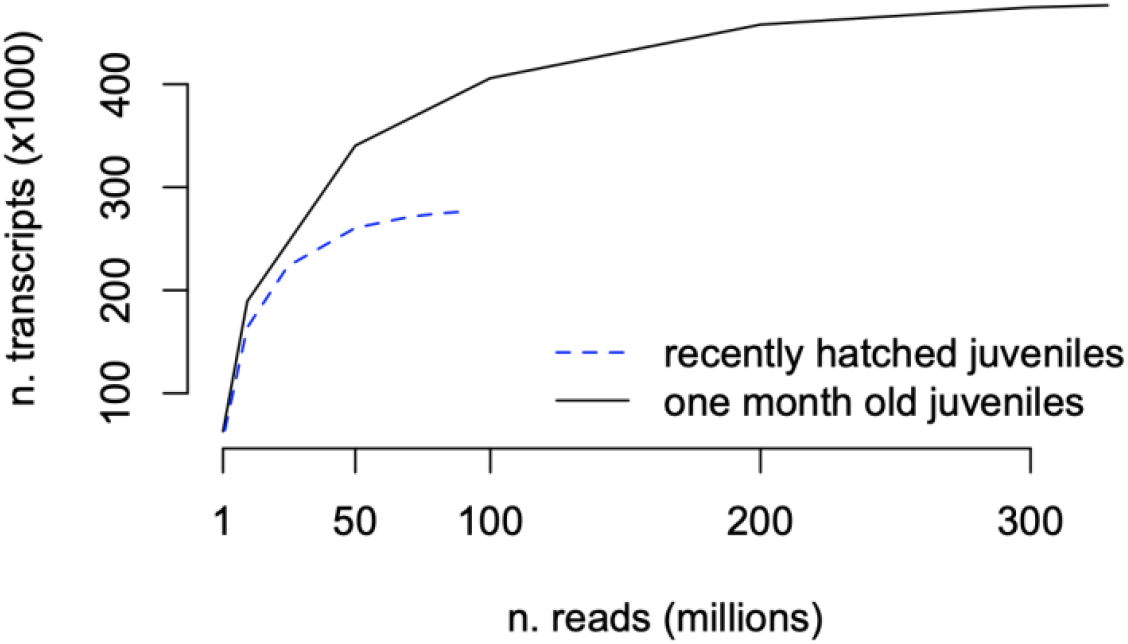
Rarefaction curves performed on the raw Trinity assemblies.

Based on this assessment, we can assume that both assemblies are of good quality in terms of transcripts length and representation of conserved transcripts.

### Transcriptome annotation

The structural annotation performed with Transdecoder on the thinned assemblies identified 42,959 and 53,992 ORFs for rnasep1 and rnasep2, respectively, which is consistent with the number of proteins reported from the reference genome (n = 50,979) (Rencken et al., 2025). After the filtering of longest ORF per transcript isoform, we obtained 35,590 ORFs for rnasep1 and 44,233 ORFs for rnasep2. These ORFs were submitted to functional annotation.

In the assembly of newly hatched individuals, 26,290 ORFs were annotated at least once. BLASTp annotated 24,299 transcripts against the SwissProt/Uniprot database (i.e. 4,958 from the curated database and 19,341 from the whole database). In addition, 21,351 orthologous hits were found by eggNOG-mapper with 14,709 transcripts annotated with gene ontology terms (for the occurrence of the top 20 GO terms from each namespace: Figure 4a). Moreover, 12,967 transcripts were associated to KEGG KO and 6,947 to KEGG pathways (Figure 5a).

**Figure 4.**
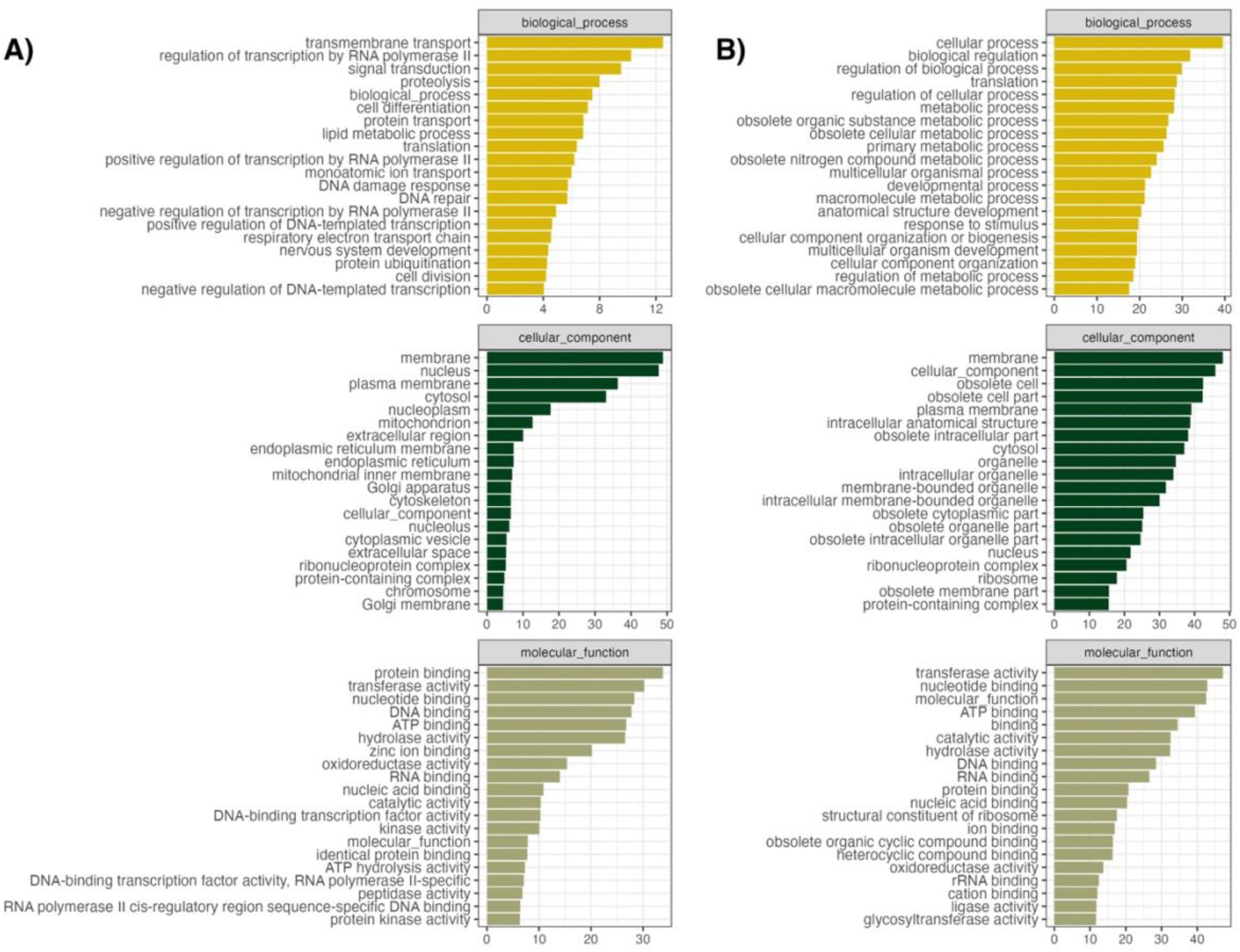
Occurrence of the 20 top GO terms related to the putative ORFs by namespace for A) newly hatched and B) one-month-old juveniles.

**Figure 5.**
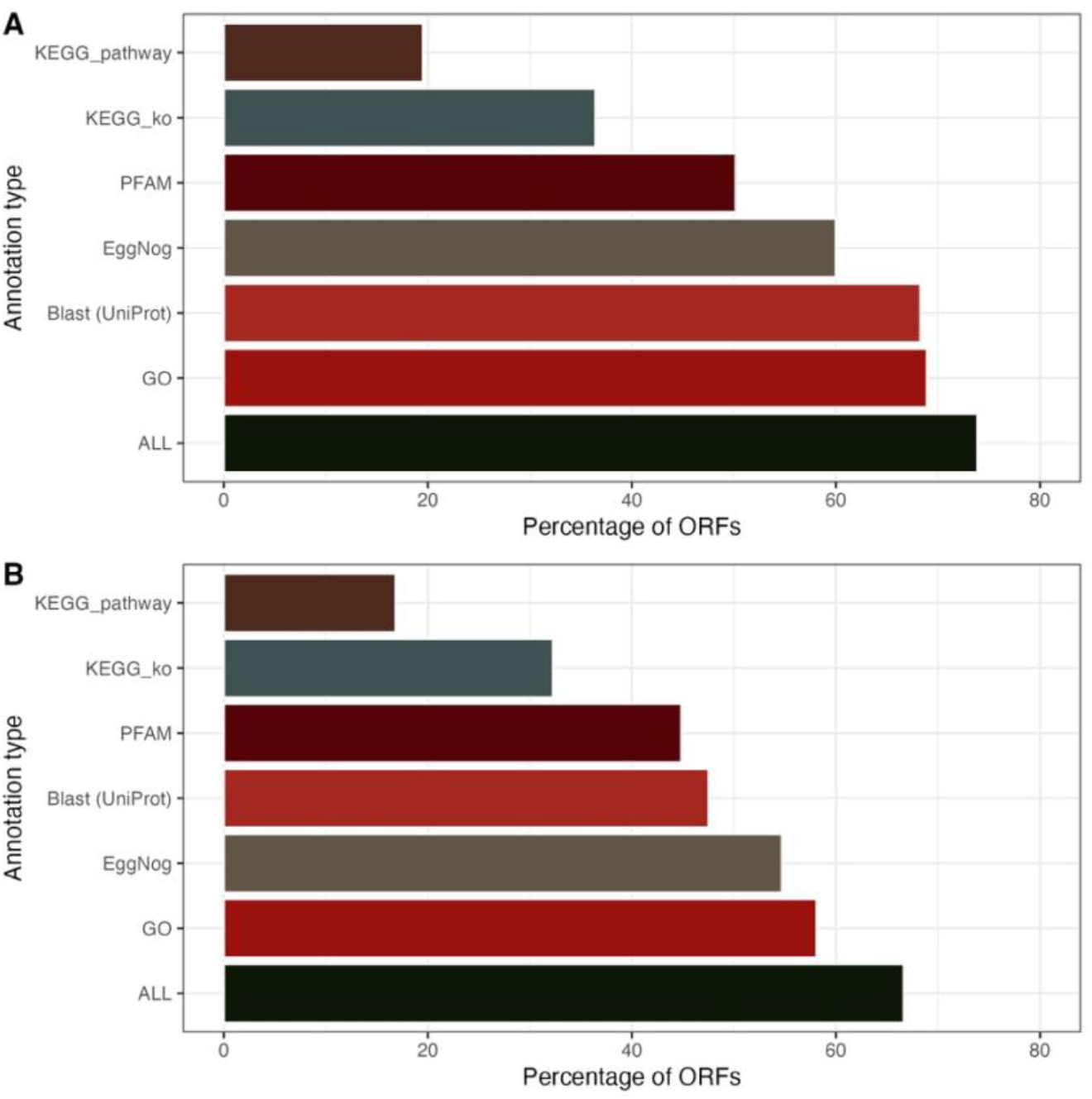
Percentage of ORFs matching at least one entry from each annotation type for A) whole body hatchlings and B) head of one-month-old juveniles.

In the one-month-old assembly, 29,474 ORFs were annotated at least once. BLASTp annotated 21,014 transcripts against the SwissProt/Uniprot database (i.e. 6,370 from the curated database and 14,644 from the whole database). This annotation was completed using EggNOG-mapper. Overall, 24,196 orthologous hits were reported and 16,204 transcripts matched gene ontology terms (Figure 4b), 14,269 transcripts matched KEGG KO and 7,444 matched KEGG pathways (Figure 5b).

### Newly hatched vs one-month-old juveniles

Overall, 17,526 transcripts were shared between rnasep1 and rnasep2 assemblies (RBH), representing 49% and 40% of each assembly, respectively (Figure 6). Consequently, 18,064 and 26,707 transcripts did not match RBH requirements in rnasep1 and rnasep2, respectively. Among them, ∼40% (rnasep1) and ∼20% (rnasep2) were classified as “putative specific” (i.e. no hit from the reciprocal BLASTp). The visualization of the top 20 unique GO categories from the “putative specific” sequences revealed consistent patterns (Figure 6). Three main functional themes are highlighted from newly hatched juveniles (whole body): early structuring of neural network (e.g. *neural crest formation, dendrite arborization* and *presynaptic modulation of chemical synaptic transmission*), response to cellular stress and protein quality (e.g. *negative regulation of PERK-mediated unfolded protein response, protein branched polyubiquitination*) and metabolism/transport/homeostasis (e.g. *glycine betaine transport, amino acid transport, plasma membrane phospholipid scrambling*). The presence of *neural crest formation* term echoes the hypothesis of cordal pattern of neurogenesis in cephalopods (Shigeno et al., 2015), and other neural structuring-related terms are consistent with the active neural development at this stage (Nixon & Mangold, 1998). The main unique GOs from one-month-old juveniles (head tissues) displayed clear pattern as well. There is a strong signal of lipid processing (e.g. *sphinganine metabolic process, unsaturated fatty acid metabolic process, acylglycerol acyl-chain remodeling*), neural signalling (e.g. *neuronal ion channel clustering, catecholamine transport, spinal cord association neuron specification*), secondary metabolism and post-translational modifications (e.g. *C-terminal protein amino acid modification, RNA 5’-end processing*).

**Figure 6.**
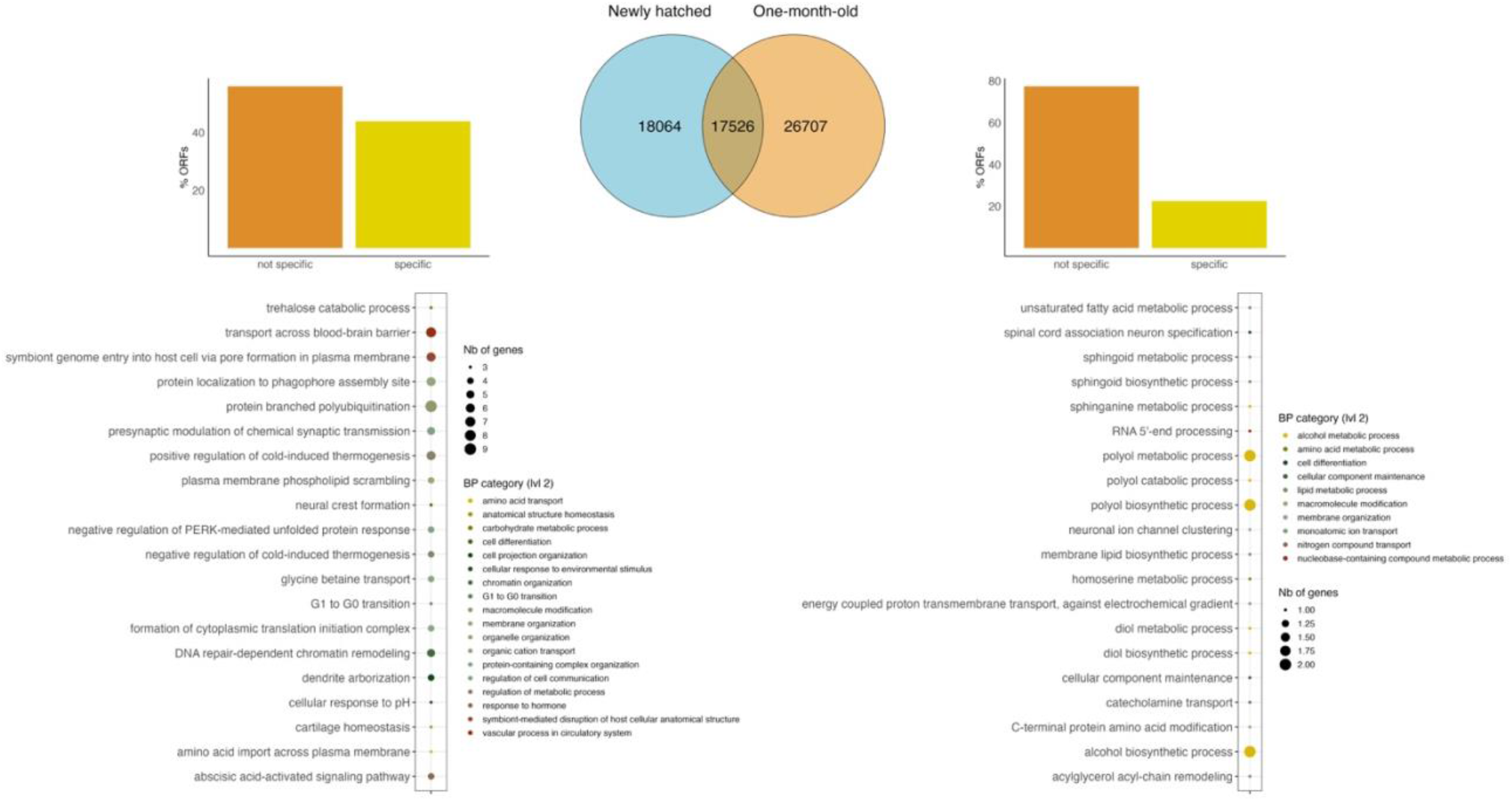
Venn diagram displaying the number of RBH and non-RBH protein sequences. The bar plots represent the proportion non-RBH sequences specific to each assembly (i.e. no Blastp hit; left: newly hatched juveniles; right: one-month-old juveniles). The top 20 (based on occurrrence) level 6 GO terms are displayed in the scatter plots (left: newly hatched juveniles; right: one-month-old juveniles).

Overall, specific transcripts from rnasep1 are dominated by developmental processes such as nervous system structuring, cellular stress control and some metabolic functions potentially implicated in the post-hatching transition (Minet et al., 2025). On the other side, specific transcripts from rnasep2 are more clearly associated with a mature neural system functioning, including lipidic remodelling of membranes, biosynthesis and metabolism of sphingolipids as well as neurotransmission. These main functions are consistent with the lipid-rich cephalopod brain (Navarro et al., 2000), which specifically contains sphingolipids (Kreps et al., 1968), and the role catecholamines (mostly dopamine and noradrenaline) play as neurotransmitter in *Sepia officinalis* nervous system (Juorio, 1971). Therefore, Figure 6 shows that specific transcripts are likely to results from biological signal related to developmental stage and tissue diversity.

In addition, the difference in the number of transcripts detected between newly hatched and one-month-old juveniles is more likely to result from biological signal than technical artefact. Indeed, from the Figure 3, we can argue that the sampling depth was enough in both assemblies to capture all the information contained in the sequencing data. Moreover, the brief GO analysis demonstrated that both assemblies encompass biological specificities resulting from the collected tissues (whole body vs head) and the developmental stage of the individuals. In this sense, cephalopod brains are known to produce a high quantity of transcripts, in particular thanks to an extraordinary rate of mRNA editing (∼60% of brain transcripts) (Rosenthal & Eisenberg, 2025; Voss & Rosenthal, 2023) resulting in a unique diversification of isoforms. And this diversification is likely to occur with the acquisition of new behaviours triggered by environmental context experienced throughout life (Fischer et al., 2021), explaining the higher number of isoforms found in one-month-old juveniles while newly hatched individuals probably express only innate behaviour-related transcripts. Moreover, rnasep1 individuals may still be under transcriptional canalization due to their temporal proximity with hatchling transition, that is embryonic stage, resulting in an overall less variable transcriptomic activity (Irvine, 2020). Finally, the predominance of brain tissues in rnasep2 while rnasep1 contains a lot of tissues producing poorly diversified transcripts (i.e. mantle, muscles) is likely to dilute signal from more plastic tissues (neural tissues), thereby lower the overall variability as well.

## Conclusion

Both transcriptome assemblies significantly contribute to improve molecular resources for conducting genomic studies on Cephalopods, particularly *S. officinalis*. The overall overlap rate (49% and 40% for rnasep1 and rnasep2, respectively) between transcripts of whole body hatchlings and head of one-month-old juveniles, as well as the functional specificity of unique sequences attest to the importance of considering developmental stages and tissues in transcriptomic studies. In this sense, the use of multiple tissues (whole body in newly hatched and heads in one-month-old juveniles), although it may obscure some tissue-specific patterns, provides valuable resources for organism-level investigations. Moreover, the inclusion of individuals from various metal and *p*CO_2_ exposure conditions diversified the transcriptomic landscapes included in the assemblies, broadening their range of relevance. This is particularly critical in the context of global change which requires to consider the influence of combined environmental stressors.

## Acknowledgement

We are grateful to the *Institut Français de Bioinformatique* and the IFB-Core cluster platform, financed under the *Programme d’Investissements d’Avenir* funded by the *Agence Nationale de la Recherche* (RENABI-IFB ANR-11-INBS-0013 and MUDIS4LS ANR-21-ESRE-0048), for providing computing resources

## Data, scripts, code and supplementary information availability

All the transcriptomic data (raw reads) are available at the European Nucleotide Archive under the accession numbers: PRJEB89715 (rnasep1) and PRJEB88225 (rnasep2), with the two assemblies as “related ENA records”: HCHA01230572 (rnasep1) and HCHB01370613 (rnasep2). The two assemblies were also deposited with their related annotation files on Zenodo with DOI: https://doi.org/10.5281/zenodo.17878811. The raw annotation files produced by the annotation process and used as input data in the R pipeline of the project are available on Zenodo with DOI: https://doi.org/10.5281/zenodo.17896603. The Github repository containing the codes used for bioinformatic analyses was released on Zenodo, accessible under the DOI: https://doi.org/10.5281/zenodo.15655834.

