## Supplemental material for "Two *de novo* transcriptome assemblies and functional annotations from juvenile cuttlefish (*Sepia officinalis*) under various metal and *p*CO2 exposure conditions"

Table S1a: Hg and Ag tissue concentrations in rnasep1 individuals (ug/g dry weight).

| Sample | Hg | Ag | Exp |
| --- | --- | --- | --- |
| CT7.7_15 | 0.029 | 0.81 | SEP1 |
| CT7.7_2 | 0.029 | 0.81 | SEP1 |
| CT7.7_93 | 0.029 | 0.81 | SEP1 |
| CT8.1_3 | 0.03 | 0.43 | SEP1 |
| CT8.1_9 | 0.03 | 0.43 | SEP1 |
| CT8.1_82 | 0.03 | 0.43 | SEP1 |
| Ag7.7_1 | NA | 4.14 | SEP1 |
| Ag7.7_2 | NA | 4.14 | SEP1 |
| Ag7.7_34 | NA | 4.14 | SEP1 |
| Ag8.1_2 | NA | 3.85 | SEP1 |
| Ag8.1_102 | NA | 3.85 | SEP1 |
| Ag8.1_27 | NA | 3.85 | SEP1 |
| Hg7.7_125 | 0.437 | NA | SEP1 |
| Hg7.7_2 | 0.437 | NA | SEP1 |
| Hg7.7_52 | 0.437 | NA | SEP1 |
| Hg8.1_36 | 0.315 | NA | SEP1 |
| Hg8.1_111 | 0.315 | NA | SEP1 |
| Hg8.1_6 | 0.315 | NA | SEP1 |

Table S1b: Hg tissue concentrations in rnasep2 individuals (ug/g dry weight).

| Sample | Hg | Exp |
| --- | --- | --- |
| CT8.1_5 | 0.07 | SEP2 |
| CT8.1_6 | 0.06 | SEP2 |
| CT8.1_38 | 0.05 | SEP2 |
| CT8.1_80 | 0.06 | SEP2 |
| CT7.5_11 | 0.07 | SEP2 |
| CT7.5_13 | 0.06 | SEP2 |
| CT7.5_46 | 0.08 | SEP2 |
| CT7.7_85 | 0.06 | SEP2 |
| Hg8.1_20 | 0.36 | SEP2 |
| Hg8.1_24 | 0.34 | SEP2 |
| Hg8.1_25 | 0.33 | SEP2 |
| Hg8.1_56 | 0.37 | SEP2 |
| Hg7.5_28 | 0.28 | SEP2 |
| Hg7.5_30 | 0.26 | SEP2 |
| Hg7.5_32 | 0.25 | SEP2 |
| Hg7.5_68 | 0.33 | SEP2 |

Table S2a: Summary of the bowtie2 alignment report for rnasep1 assembly

| Sample | AlignmentRate (%) | AlignedOnce (%) | MultiAligned(%) | AlignedDiscordantly (%) | NotAligned (%) | Assembly |
| --- | --- | --- | --- | --- | --- | --- |
| Ag7.7_1 | 90.7 | 50.84 | 34.97 | 4.89 | 9.3 | thinned |
| Ag7.7_2 | 91.3 | 49.05 | 37.88 | 4.37 | 8.7 | thinned |
| Ag7.7_34 | 91.68 | 48.89 | 37.81 | 4.98 | 8.32 | thinned |
| Ag8.1_102 | 91.96 | 48.95 | 37.72 | 5.29 | 8.04 | thinned |
| Ag8.1_27 | 91.38 | 49.08 | 37.03 | 5.27 | 8.62 | thinned |
| Ag8.1_2 | 90.36 | 49.52 | 36.36 | 4.48 | 9.64 | thinned |
| CT7.7_15 | 91.26 | 49.38 | 36.58 | 5.3 | 8.74 | thinned |
| CT7.7_2 | 90.6 | 50.15 | 35.21 | 5.24 | 9.40 | thinned |
| CT7.7_93 | 91.28 | 48.55 | 37.85 | 4.88 | 8.72 | thinned |
| CT8.1_3 | 91.59 | 50.53 | 35.72 | 5.34 | 8.41 | thinned |
| CT8.1_82 | 91.91 | 48.17 | 38.36 | 5.38 | 8.09 | thinned |
| CT8.1_9 | 91.34 | 50.03 | 35.47 | 5.84 | 8.66 | thinned |
| Hg7.7_125 | 91.39 | 49.05 | 36.46 | 5.88 | 8.61 | thinned |
| Hg7.7_2 | 91.12 | 47.94 | 36.75 | 6.43 | 8.88 | thinned |
| Hg7.7_52 | 91.45 | 48.99 | 35.92 | 6.54 | 8.55 | thinned |
| Hg8.1_111 | 92.26 | 48.9 | 36.99 | 6.37 | 7.74 | thinned |
| Hg8.1_36 | 91.32 | 48.48 | 37.91 | 4.93 | 8.68 | thinned |
| Hg8.1_6 | 91.24 | 49.19 | 37.04 | 5.01 | 8.76 | thinned |
| Ag7.7_1 | 97.83 | 41.44 | 51.81 | 4.58 | 2.17 | raw |
| Ag7.7_2 | 97.86 | 40.97 | 52.8 | 4.09 | 2.14 | raw |
| Ag7.7_34 | 97.85 | 38.45 | 54.64 | 4.76 | 2.15 | raw |
| Ag8.1_102 | 97.87 | 39.99 | 52.86 | 5.02 | 2.13 | raw |
| Ag8.1_27 | 97.74 | 40.51 | 52.2 | 5.03 | 2.26 | raw |
| Ag8.1_2 | 97.86 | 40.48 | 53.21 | 4.17 | 2.14 | raw |
| CT7.7_15 | 97.75 | 40.49 | 52.21 | 5.05 | 2.25 | raw |
| CT7.7_2 | 97.62 | 40.18 | 52.47 | 4.97 | 2.38 | raw |
| CT7.7_93 | 97.73 | 40.75 | 52.37 | 4.61 | 2.27 | raw |
| CT8.1_3 | 97.99 | 40.59 | 52.38 | 5.02 | 2.01 | raw |
| CT8.1_82 | 97.84 | 39.19 | 53.54 | 5.11 | 2.16 | raw |
| CT8.1_9 | 97.82 | 40.3 | 52.01 | 5.51 | 2.18 | raw |
| Hg7.7_125 | 97.62 | 40.27 | 51.71 | 5.64 | 2.38 | raw |
| Hg7.7_2 | 97.92 | 37.86 | 53.76 | 6.3 | 2.08 | raw |
| Hg7.7_52 | 97.74 | 38.25 | 53.18 | 6.31 | 2.26 | raw |
| Hg8.1_111 | 97.93 | 36.78 | 55.16 | 5.99 | 2.07 | raw |
| Hg8.1_36 | 97.76 | 40.45 | 52.65 | 4.66 | 2.24 | raw |
| Hg8.1_6 | 97.88 | 40.95 | 52.14 | 4.79 | 2.12 | raw |

Table S2b: Summary of the bowtie2 alignment report for rnasep2 assembly.

| Sample | AlignmentRate (%) | AlignedOnce (%) | MultiAligned (%) | AlignedDiscordantly (%) | NotAligned (%) | assembly |
| --- | --- | --- | --- | --- | --- | --- |
| CT8.1_85 | 93.37 | 46.65 | 43.71 | 3.01 | 6.63 | thinned |
| Hg7.7_20 | 93.76 | 43.6 | 45.71 | 4.45 | 6.24 | thinned |
| Hg7.7_24 | 93.86 | 48.69 | 43.34 | 1.83 | 6.14 | thinned |
| Hg7.7_25 | 93.79 | 47.28 | 43.86 | 2.65 | 6.21 | thinned |
| Hg7.7_56 | 92.99 | 46.62 | 43.11 | 3.26 | 7.01 | thinned |
| Hg8.1_28 | 92.88 | 45.18 | 43.29 | 4.41 | 7.12 | thinned |
| Hg8.1_30 | 93.34 | 47.33 | 42.77 | 3.24 | 6.66 | thinned |
| Hg8.1_32 | 92.32 | 45.77 | 41.72 | 4.83 | 7.68 | thinned |
| Hg8.1_68 | 92.28 | 49.01 | 40.99 | 2.28 | 7.72 | thinned |
| CT8.1_13 | 93.91 | 45.88 | 45.68 | 2.35 | 6.09 | thinned |
| CT8.1_46 | 92.94 | 46.37 | 43.69 | 2.88 | 7.06 | thinned |
| CT7.7_80 | 93.04 | 47.18 | 43.34 | 2.52 | 6.96 | thinned |
| CT7.7_6 | 94.03 | 45.61 | 45.63 | 2.79 | 5.97 | thinned |
| CT8.1_11 | 94.00 | 47.06 | 44.32 | 2.62 | 6.00 | thinned |
| CT7.7_5 | 93.54 | 46.54 | 43.38 | 3.62 | 6.46 | thinned |
| CT7.7_38 | 93.74 | 46.93 | 43.97 | 2.84 | 6.26 | thinned |
| CT8.1_85 | 98.75 | 33.23 | 62.7 | 2.82 | 1.25 | raw |
| Hg7.7_20 | 98.63 | 31.36 | 63.17 | 4.1 | 1.37 | raw |
| Hg7.7_24 | 98.71 | 33.64 | 63.42 | 1.65 | 1.29 | raw |
| Hg7.7_25 | 98.52 | 32.36 | 63.67 | 2.49 | 1.48 | raw |
| Hg7.7_56 | 98.43 | 33.36 | 62.00 | 3.07 | 1.57 | raw |
| Hg8.1_28 | 97.87 | 31.24 | 62.29 | 4.34 | 2.13 | raw |
| Hg8.1_30 | 98.41 | 32.2 | 63.19 | 3.02 | 1.59 | raw |
| Hg8.1_32 | 98.16 | 33.8 | 59.79 | 4.57 | 1.84 | raw |
| Hg8.1_68 | 98.64 | 36.74 | 59.91 | 1.99 | 1.36 | raw |
| CT8.1_13 | 98.89 | 32.2 | 64.54 | 2.15 | 1.11 | raw |
| CT8.1_46 | 98.89 | 34.51 | 61.83 | 2.55 | 1.11 | raw |
| CT7.7_80 | 98.76 | 34.49 | 61.98 | 2.29 | 1.24 | raw |
| CT7.7_6 | 98.85 | 31.71 | 64.63 | 2.51 | 1.15 | raw |
| CT8.1_11 | 98.97 | 33.27 | 63.38 | 2.32 | 1.03 | raw |
| CT7.7_5 | 98.67 | 32.58 | 62.7 | 3.39 | 1.33 | raw |
| CT7.7_38 | 98.97 | 33.51 | 62.94 | 2.52 | 1.03 | raw |
